# Fusing spheroids to aligned μ-tissues in a Heart-on-Chip featuring oxygen sensing and electrical pacing capabilities

**DOI:** 10.1101/2022.02.26.482011

**Authors:** Oliver Schneider, Alessia Moruzzi, Stefanie Fuchs, Alina Grobel, Henrike S. Schulze, Torsten Mayr, Peter Loskill

## Abstract

Over the last decade Organ-on-Chip (OOC) emerged as a promising technology for advanced *in vitro* models, recapitulating key physiological cues. OOC approaches tailored for cardiac tissue engineering resulted in a variety of platforms, some of which integrate stimulation or probing capabilities. Due to manual handling processes, however, a large-scale standardized and robust tissue generation, applicable in an industrial setting, is still out of reach. Here, we present a novel cell injection and tissue generation concept relying on spheroids, which can be produced in large quantities and uniform size from induced pluripotent stem cell-derived human cardiomyocytes. Hydrostatic flow transports and accumulates spheroids in dogbone-shaped cultivation chambers, which subsequently fuse and form aligned, contracting cardiac muscle fibers. Furthermore, we demonstrate electrical stimulation capabilities by utilizing fluidic media connectors as electrodes and provide the blueprint of a low-cost, open-source, scriptable pulse generator. We report on a novel integration strategy of optical O_2_ sensor spots into resin-based microfluidic systems, enabling *in situ* determination of O_2_ partial pressures. Finally, proof-of-concept demonstrating electrical stimulation combined with *in situ* monitoring of metabolic activity in cardiac tissues is provided. The developed system thus opens the door for advanced OOCs integrating biophysical stimulation as well as probing capabilities and serves as blueprint for the facile and robust generation of high density microtissues in microfluidic modules amenable for scale-up and automation.

## 1. Introduction

Microfluidic Organ-on-Chip (OOC) systems have emerged over the last decade as auspicious alternative to conventional cell culture methods, enabling the generation of tissues in precisely controlled environments featuring vasculature-like perfusion. Numerous OOCs culturing single tissues or even combinations of multiple organs have been presented as well as their applicability for mechanistic biomedical research, personalized medicine and drug testing.

Since cardiotoxicity plays a major role not only in the drug development process, various Heart-on-Chip (HoC) platforms mimicking the native myocardium, i.e., generating an aligned cardiac muscle fiber have been developed. [1–4] However, the bare generation of physiologically more relevant model systems is only a first step in investigating and understanding human pathophysiology of specific diseases. For gaining advanced insights, it is just as well important to study the effects of precisely controlled external stimuli, and, to record tissue-specific key parameters. [5] Therefore, one major goal in OOC technology is a direct chip integration of mentioned capabilities.

Due to the unique excitability of cardiac tissue, one of the most important stimuli is electrical stimulation. Pacing can, i.a., be used for advanced tissue maturation or for synchronization of tissue beating, yielding a defined baseline for controlled drug testing. [3,6] Current microphysiological systems integrate pacing capabilities with varying degrees of fabrication complexity, e.g., by utilizing substrate-integrated thin film electrodes, integrating wires into the chip, or fabricating conductive PDMS pillars. [7–9] In addition to electrical stimulation, mechanical, chemical, or combined stimulations have been presented. [10–12]

Similarly, various approaches have been pursued for directly integrating sensing capabilities into HoCs, offering insights into electrophysiology, barrier integrity, or O_2_ levels. [13–16] As O_2_ plays a key role in regulating cell functions and O_2_ consumption can provide crucial insights into cell state and tissue metabolism, the *in situ* determination of O_2_ concentration is of particular interest. More broadly, a simple sensor integration is not only of relevance for O_2_ monitoring or for cardiac OoCs in general but universally desired for all tissue types. For this purpose, luminescent optical sensor spots are an elegant approach as they can be integrated into microphysiological systems without the need of a wired chip connection, offering on-demand readout opportunities. [17–19]

For an accurate recapitulation of native cardiac tissue, high cell densities leading to an abundance of cell-cell contacts within cultured tissue construct are crucial. Due to the post-mitotic nature of cardiomyocytes (CMs), the required cell density has to be achieved already during the initial cell injection process. Nevertheless, many systems currently rely on the injection of cell-laden hydrogels, yielding low cell densities. On the other hand, flow-based injection of cell suspensions usually deploys micron-sized retaining features for accumulating cells in defined geometries, which, however, are prone to clogging during the loading process, yielding uncontrolled loading pressures, ceasing of flow and low densities.

As promising alternative, we recently presented centrifugation-based cell injection techniques. [20,21] Generally, however, the injection of single cell suspensions strongly relies on a well singularized suspension and is prone to inhomogeneities, lacking robustness due to naturally occurring polydispersity. Contrary, spheroids of CMs can be generated with precisely controlled dimensions and have been shown to be able to fuse to arbitrary tissue shapes or be guided inside microfluidic environments. [22–24]

Here, we present a novel OOC loading and tissue generation concept based on the introduction of preformed spheroids of controlled size into dogbone-shaped tissue chambers via a facile and robust hydrostatic loading. The individual spheroids, formed from human induced pluripotent stem cell (hiPSC) derived CMs subsequently merged to an aligned beating cardiac µ-tissue inside the microfluidic system.

Moreover, we demonstrate the integration of both electrical pacing as well as O_2_ readout capabilities into the HoC. By using existing fluidic media connectors as electrodes, we confirm reliable tissue excitation with minimal changes in the experimental setup, without the need for advanced fabrication techniques. We report on a newly developed, remotely controllable, “do-it-yourself (DIY)” pacemaker, a valuable alternative to specialized cost-intensive experimental hardware. Furthermore, we introduce a novel strategy for integrating luminescent optical sensor spots into microphysiological platforms by assembling the chip directly from photocurable polymer on top of a sensor substrate. By ultimately combining stimulation and probing capabilities, we reveal changes in O_2_ consumption of cardiac tissues upon varying electrical stimulation inside the HoC.

## 2. Materials and methods

### 2.1. Chip fabrication

The HoC is composed of a polyethylene-terephthalate (PET) substrate with deposited O_2_ sensors, a tissue stencil molded from UV curable resin (NOA 81, Norland adhesive, USA), a media module out of polydimethylsiloxane (PDMS) and a track-etched PET membrane with a pore size of 3 µm (03044, SABEU, Germany), which was plasma coated following previously published procedure. [25] The media module and tissue layer patterning stamp were manufactured by soft lithography and replica molding with PDMS. [26] Briefly, SU-8 based (SU-8 50 & SU-8 100, MicroChem, USA) wafer masters were fabricated according to the manufacturer’s protocols (media module: positive structures, 150 µm channel, SU8-100; tissue stamp: negative structures, 150 µm channel, SU-8 100 and 30 µm constriction, SU8-50). Trichloro(1H,1H,2H,2H-perfluorooctyl)silane (448931, Sigma-Aldrich, USA) was vapor deposited on both masters for 1 h to facilitate demolding of PDMS. PDMS (Sylgard 184, Dow Corning, USA) was mixed (10 : 1 base to curing agent mass ratio), degassed and poured onto the wafers. After overnight curing at 60 °C, 3 mm thick PDMS replicates were peeled off the wafer and cropped to chip size. Inlets for media and tissue channels were punched with a biopsy puncher (504529, World Precision Instruments, Germany) into the media module. Access ports for resin injection were punched with a biopsy puncher (504531, World Precision Instruments, Germany) in all four corners of the injection cavity region of the tissue patterning stamp.

PDMS replicas of the media layer were cleaned with isopropanol (IPA) and residual particles removed using adhesive tape (56002-00001-01, Tixo, Austria). PET-membranes were cut to desired size using a CO_2_ laser cutter (VLS2.30, Universal Laser Systems, USA) and rinsed in ethanol. Following O_2_ plasma treatment of both parts (15 s, 50 W; Zepto, Diener, Germany), the media module was aligned onto the membrane, covering both chambers. The assembly was baked at 60 °C for 2 h and stored until further usage.

The oxygen indicator dye platinum(II)meso-tetra(4-fluorophenyl) tetrabenzoporphyrin (Pt-TPTBPF) was synthesized as previously described. [27] A stock solution of 10 % polystyrene (260 kDa) in Toluene was prepared. 1 mg oxygen indicator dye was added to 1 g stock solution yielding a concentration of 1 % dye in polymer after solvent evaporation. The solution was homogenized by vigorous stirring for 10 minutes.

PET films (Melinex® 506, DuPont Teijin Films, USA) with a thickness of 125 µm were used as sensor substrate. The sensor solution was applied using a microdispenser (MDS3200+, VERMES Microdispensing GmbH, Germany) equipped with a 70 µm nozzle and tungsten tappet with a tip diameter of 0.7 mm. The microdispenser was mounted on a custom-made CNC platform for precise positioning of sensor spots. Detailed printing parameters are listed in Table T1.

The sensor substrate was rinsed with IPA, blow dried, treated with O_2_ plasma (60 s, 50 W; Zepto, Diener, Germany) and covered with a solution of 1 % (3-Aminopropyl)triethoxysilan (APTES, A3648, Sigma-Aldrich, USA) dissolved in DI water. After 10 min the sensor substrate was rinsed with deionized (DI) water and blow dried. The tissue stamp was cleaned with IPA and adhesive tape and aligned onto the sensor substrate, ensuring full contact of stamp structures with the substrate while avoiding collapsing of the glue injection cavity. NOA 81 was inserted into a 10 ml syringe (BD Plastipak, BD, USA) with attached dispensing tip (21 GA; KDS212P, Weller, USA) and both resin injection ports were filled to the top. Resin started to immediately fill the microcavity. For complete molding, injection ports were filled again after a stopping of filling was observed. After 20 min the molding cavity was completely filled and the assembly exposed to UV light at 188 mJ/cm^2^ (λ = 365 nm; LED UV mini-Oven, Novachem, Ireland). The stamp was carefully removed, cleaned with adhesive tape and stored for repeated use. Prepared media-membrane assembly was cleaned with IPA and adhesive tape (area not covered with membrane). Following plasma treatment (60 s, 50 W; Zepto, Diener, Germany), the surface was covered with a 1 % APTES solution in DI water for 10 min. The module was rinsed in DI water, blow dried and aligned onto the tissue stencil molded from resin. Slight pressure was applied to remove trapped air bubbles and the assembly was cured with UV light at 21 J/cm^2^ (λ = 365 nm; LED UV mini-Oven, Novachem, Ireland). Finalized chip was baked for 12 h at 60 °C to improve adhesion. Channels were rinsed with ethanol to remove any remaining, potentially cytotoxic, glue components and left to dry.

### 2.2. Cell culture

#### 2.2.1. hiPSC culture and CM differentiation

CMs were differentiated from the hiPSC line, Coriell GM25256 (RRID: CVCL_Y803, Gladstone Institute for Cardiovascular Disease, San Francisco, USA). After thawing, cells were plated on growth factor-reduced Matrigel (354277, Corning, USA)-coated 6-well plates at a density of 25000 cells/cm^2^. Cells were cultured in TeSR-E8 (05990, STEMCELL Technologies, Canada) medium, supplemented with 10 μM ROCK inhibitor Y-27632 (RI; 05990, STEMCELL Technologies, Canada) for the first 24 h after thawing or passaging. hiPSCs were passaged with Accumax (SCR006, Sigma-Aldrich, USA) at least once before initiation of differentiation.

Differentiation was achieved using an optimized protocol for the small-molecule manipulation of Wnt signaling adapted from Lian et al. [28] Upon reaching ≥ 90 % confluence (day 0) media was exchanged to RPMI 1640 medium (RPMI; 1185063, Gibco, USA) supplemented with B27 supplement without insulin (B27-I; A1895601, Gibco, USA) and 10 μM of the Wnt agonist CHIR99021 (CHIR; S2924, Selleckchem, USA). Exactly after 24 h, medium was changed to RPMI + B27-I. After 2 days (day 3), the medium was changed to RPMI + B27-I with 5 μM Wnt inhibitor IWP-4 (SML1114, Sigma-Aldrich, USA) and incubated for 48 hours. On day 5, medium was changed to RPMI + B27-I and on day 7 to RPMI 1640 supplemented with B27 complete supplement (B27C; 17504044, Gibco, USA), which was thereafter used for CM culture, and exchanged every second day. At around day 9-12, cells showed spontaneous beating and were dissociated on day 15 by incubation with 280 U/ml Collagenase (LS004174, Worthington, USA) and 40 U/ml DNase (LS006331, Worthington, USA) in RPMI + B27C for 1.5 h. Detached monolayers were collected in a 50 ml conical centrifuge tube containing 20 ml of PBS-, centrifuged at 200 x g for 3 min and resuspended in Accumax solution followed by 25 min of incubation at 37 °C. After singularization, cells were washed and resuspended in RPMI + B27C with 10 µM of RI. At this point, CM purity was checked by flow cytometry (Guava® easyCyte 8HT, Merck Millipore, USA) for the cardiac marker Cardiac troponin T conjugated to APC (cTnT; 1:50; REA400; 130-120-543, Miltenyi Biotec, Germany), and cells were frozen in RPMI + B27C with 10 µM of RI + 10% FCS + 10% DMSO. Only hiPSC-derived CMs with purity ≥ 80% were used for further experiments.

Before loading, CMs were thawed in RPMI + B27C with 10 µM of RI and plated at a cell density of 2 Mio cells/well in a 6-well plate coated with Matrigel. After 24 h media was exchanged to RPMI + B27C and after 3 days of culture, cells were used for experiments.

#### 2.2.2. Primary human dermal fibroblasts

Primary human dermal fibroblasts (phDF) were isolated from juvenile foreskin and expanded in culture. 1 Mio cells were plated in a T175 flask in DMEM (P04-04515, Pan-Biotech, Germany) supplemented with 10% FCS and 1% P/S (DMEM complete). Media was changed every 3 days. After 7 days, the cells were confluent and were either passaged or cryopreserved in DMEM media + 10% FCS + 10% DMSO. Cells were passaged at least once before any experiment and only used for experiments up to passage 9. PhDF were analyzed by flow cytometry using CD90 antibody conjugated to APC-Vio770 (CD90; 1:50; REA897; 130-114-905, Miltenyi Biotec, Germany).

#### 2.2.3. Spheroid formation

Spheroids were formed following a previously established approach. [29] Briefly, wells of a 6-well microwell culture plate (AggreWell™400, STEMCELL Technologies, USA) were replicated out of Hydrosil (101301, SILADENT, Germany). [30] Both silicone components were mixed (1:1 wt), 2.5 g was added into each well and the well plate was centrifuged at 55 x g for 60 s. After curing the silicone for 1 h at 60 °C, well replicas were carefully removed, and circular segments punched out (d = 20 mm). Circular segments were glued with an epoxy adhesive (UHU PLUS sofortfest, UHU, Germany) to a PMMA holder such that the structured side represented the bottom surface of a well (d = 15.5 mm, h = 2 mm) which could be remolded with agarose, fitting into a well of a 24-well plate.

This reusable master mold was sterilized with 70 % ethanol before every experiment. 650 µl of 3 % agarose solution in DMEM, liquified by preheating in a microwave, was deposited onto the master mold and solidified within 10 min. Once solid, agarose molds were inserted into wells of a 24-well plate with the structured side pointing upwards. 1 ml/well of PBS- was then added to each well and the plate centrifuged at 1300 x g for 3 min to remove any air bubbles trapped at the bottom of the inverted pyramid microstructure.

CMs and phDFs were dissociated by incubation with 0.05% Trypsin/Versene for 10 minutes at RT. Wells were flushed with respective media and cells transferred into a 15 ml conical centrifuge tube and centrifuged for 3 min at 200 x g. CMs were resuspended in RPMI + B27C containing 10 µM of RI, phDF in DMEM complete. Cells were mixed at a ratio of 3:1 (CMs : phDFs). A total of 0.5 Mio cells was added to each well and centrifuged at 300 x g for 3 min with a deceleration ramp setting of 3. Cells were incubated (37 °C, 5 % CO_2_) for 24 h. The next day, around 1200 spheroids/well were formed and could be used for chip loading.

#### 2.2.4. Loading protocol

Chips were treated with O_2_ plasma (60 s, 50 W; Zepto, Diener, Germany) for sterilization and hydrophilization and subsequently either filled for coating with a fibronectin solution (20 µg/ml in PBS-; F1141, Sigma-Aldrich, USA) or with pre-warmed media. Coated chips were incubated for 2 h (37 °C). Pipet tips with 100 µl media were inserted into all tissue and media access ports and remaining air bubbles removed by manual flushing. Spheroids were removed from the well, collected in a 50 ml conical centrifuge tube and sedimented by gravity. Supernatant was aspirated and the spheroids were resuspended in 250 µl/well of RPMI + B27C. 100 µl media were added to the tissue inlet tip followed by 30 µl of spheroid suspension. The difference in liquid column height led to a hydrostatic flow dragging spheroids within several minutes into the cultivation chamber, where they accumulated due to the constricting channel height. The loading process was monitored with a microscope, such that additional spheroids could be added to the inlet tip if the injected amount of spheroids was not sufficient.

#### 2.2.5. Chip culture

Following satisfactory loading, media inlet and outlet tips were topped with additional 150 µl of media and the chips were placed in a CO_2_-controlled incubator (5 % CO_2_, 37 °C, 95 % relative humidity) overnight. The next day, pipet tips were removed from the tissue channel and corresponding ports sealed with stainless steel plugs. The media channel was subsequently perfused with media at a constant flow rate of 50 μl/h using an external syringe pump (LA-190, Landgraf HLL, Germany). For pacing and O_2_ monitoring experiments, chips were removed with corresponding syringes from the incubator, placed into a cell culture hood with incubation conditions (5 % CO_2_, 37 °C, 72 % relative humidity; Incubator FlowBox™, ALS, Germany) and reattached to the syringe pump.

### 2.3. Immunofluorescence staining

Tissues were fixed inside the chips at room temperature for 15 min by inserting a 4 % solution of Roti®Histofix (P087, Carl Roth, Germany), and then permeabilized for 15 min with 0.1 % Triton X-100 (28314, Thermo Fisher Scientific, USA). The medium layer was removed, facilitated by a weaker bonding of the membrane-covered region between tissue and media module. After blocking for 1 h by adding a drop of 3 % bovine serum albumin (A9418, Sigma-Aldrich, USA) onto exposed tissue, the tissue was incubated with added staining buffer overnight at 4 °C. The staining buffer was composed of APC conjungated cTnT antibody (final dilution: 1:50), DAPI (1 mg/mL, final dilution: 1:250; D9542 Sigma-Aldrich, USA), 0.1 % Saponin (84510-100G, Sigma-Aldrich, USA), and 0.1 % bovine serum albumin (A9418, Sigma-Aldrich, USA), diluted in PBS^-^. Between steps, tissues were flushed with PBS^-^. For imaging, the chip was flipped onto a #1.5 coverslip and imaged with a magnification of up to 63 x using a laser scanning microscope (LSM 710; Zeiss, Germany).

### 2.4. Characterization of beating motion

Contraction and relaxation motion was analyzed in recorded videos of beating cardiac tissues with OpenHeartWare v1.3 (https://github.com/loslab/ohw) using the optical flow algorithm with a block width of 16 pixels, delay of 2 frames and maximum shift of 7 px. For 1D representations of beating kinetics, mean absolute motion was calculated from obtained vector field. Peak positions were automatically detected and corresponding beating frequencies determined.

### 2.5. Simulation of electrical field distribution

Electrical field strength within the paced chip was simulated using finite elements (COMSOL Multiphysics® v5.6, COMSOL AB, Sweden) in a stationary study with the electric currents physics interface. Electrodes were modeled as cylinders in media inlet and outlet with a potential difference of U = 10 V between the surfaces of both electrodes. PDMS was treated as insulator and a conductivity of σ = 1.5 S/m and relative permittivity of ε_r_ = 80.1 was assigned to the media domain, adapting previously published material parameters. [8] The absolute field strength was calculated from individual spatial components and plotted in the vertical midplane of the media channel (z = 305 µm) and the tissue chamber (z = 90 µm).

### 2.6. Pacing of tissues

Tissues were field-paced using the custom-built Arduino-based electrical stimulator “Easypace”. The stimulator was able to deliver biphasic pulses (U = -U_0_ for t = t_w_, followed by U = +U_0_ for t = t_w_, repeated after t = 1/f) in adjustable 0.1 Hz frequency steps, 1 ms pulse width steps at an amplitude of up to U_0_ = 12. 5 V and allowed a scripted control of pacing parameters. We provide extensive build-instructions for this cost-efficient (< 100 €) open-source tool (https://github.com/loslab/easypace/). If not stated otherwise, a pulse width of t_w_ = 50 ms and amplitude of U_0_ = 10 V were used, these parameters were determined as most robust for inducing pacing in investigated systems.

### 2.7. Optical analysis of O_**2**_

Chips were fixed with magnets on a custom-built sensor platform out of PMMA containing cutouts with geometrically matching sensor positions of four adjoining chips. Optical fibers could be press fitted into provided cutouts, enabling a flexible configuration of the measurement setup, e.g., measuring signals of both sensor spots of two chips, or measuring signals of one sensor spot per chip of four chips in total. Sensors were read out using a 4-channel phasefluorimeter (FireSting pro, Pyroscience, Germany) with connected optical fibers (d = 1 mm, l = 1 m). The phasefluorimeter was set to an illumination intensity of 100 % and detection amplification of 400 x. Chips were manually aligned for an optimal positioning of the sensor spot above the fiber, yielding signal intensities above 100 mV for every measurement. All measurements were carried out with the same phase shift calibration, previously obtained from a phase shift measurement of the bare sensor substrate at 37 °C, covered with well-aerated water (pO_2_ = 100 %, Δϕ = 20.4°) or a 10 % Na_2_SO_3_ solution (pO_2_ = 0 %, Δϕ = 53.8°).

## 3. Results

### 3.1. Concept of the Spheroflow HoC system

#### 3.1.1. Tissue generation

The underlying tissue generation concept relies on an initial formation of multicellular 3D cardiac µ-tissues. (Fig. 1 A) Spheroid formation is a widely established technique yielding 3D tissues without the use of hydrogel. While achieving physiological densities and cell-cell contacts, they cannot recapitulate structural organization and cell alignment, recapitulating human physiology only partially. The Spheroflow HoC concept is based on injection of pre-formed spheroids in a dogbone-shaped geometry leading to a subsequent fusion of the individual spheroids to an uniaxially aligned cardiac muscle fiber. Generation of spheroids enables a precisely controlled cellular tissue composition by mixing defined ratios of different cell types, such as CMs and fibroblasts (FBs).

**Figure 1:**
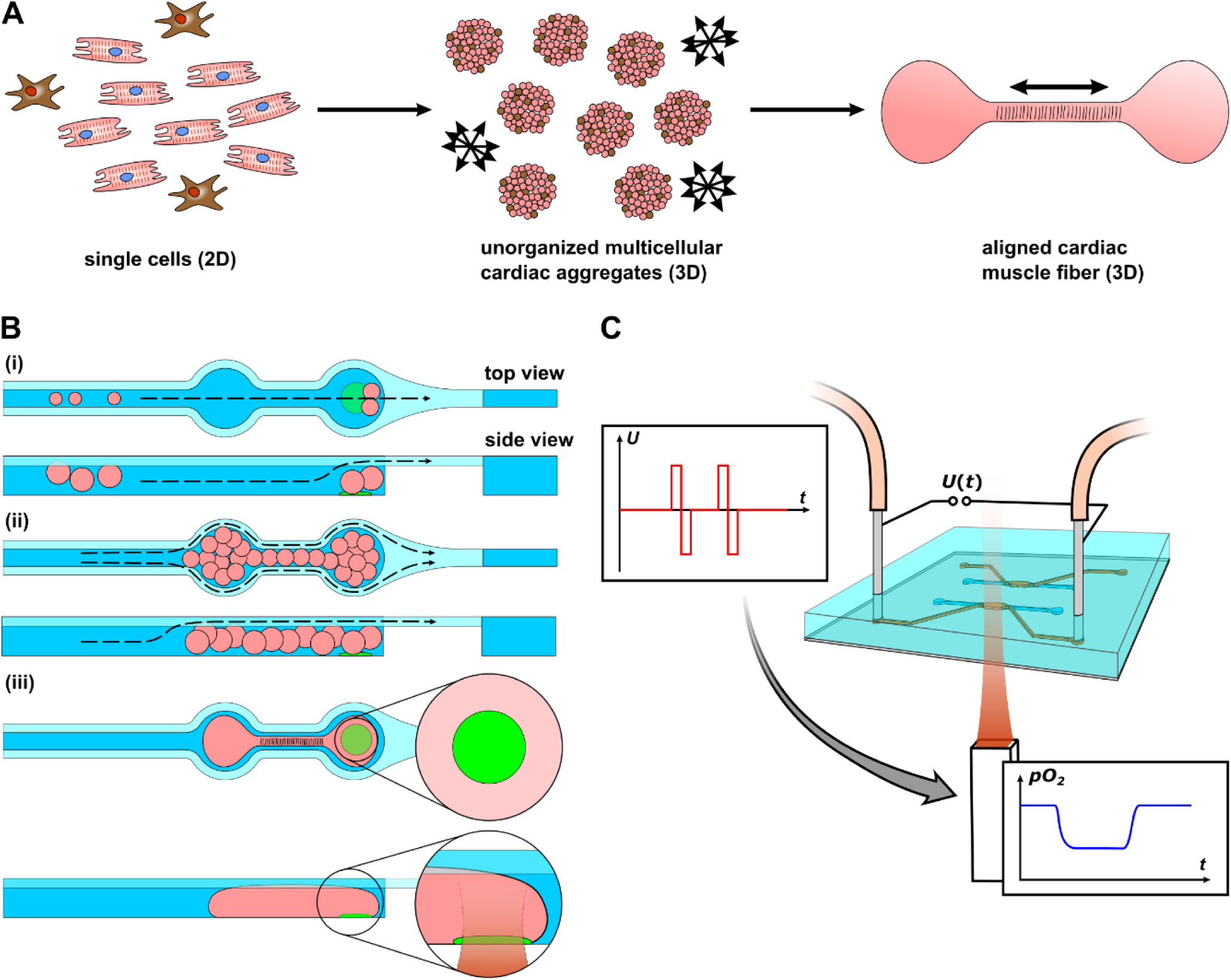
Concept of the Spheroflow HoC: A) Schematic of the bottom-up approach for generating aligned cardiac fibers. Initial compaction of single cells of precisely defined mixture to 3D spheroids. Spheroids further merge to an aligned tissue fiber, guided by provided chamber shape. Double headed arrows indicate alignment direction. B) Chip loading mechanism. (i) Spheroids are introduced by hydrostatically driven flow into the dogbone-shaped tissue compartment and fixed by the channel constriction. (ii) With gradual filling, the loading flow is maintained by the side lying constriction channel. (iii) Individual spheroids merge to a single aligned cardiac tissue. Embedded O_2_ sensor in the knob region enables in situ readout of O_2_ levels C) Investigation of metabolic activity in perfused tissues. Presented platform enables electrical tissue stimulation coupled to optical readout of O_2_ levels.

#### 3.1.2. Loading mechanism

The design of the tissue module enables the injection of the spheroids simply by hydrostatic pressure driven flow. The module is composed of structures featuring two heights, namely a channel of 180 µm ending in a dogbone-shaped tissue chamber and a channel with a height of 30 µm. Spheroids are loaded into the tissue chamber by adding the spheroid suspension into a pipet tip in the inlet. The difference in liquid column height, with respect to the tissue outlet, induces a hydrostatic flow, dragging spheroids into the channel. (Fig. 1B i) As the channel constricts at the end of the tissue chamber to the narrow channel height, spheroids are not flushed to the outlet and accumulate inside the tissue chamber. (Fig. 1 B ii) Clogging of the channel is avoided by a narrow side channel into which spheroids cannot penetrate, such that a constant flow is maintained during filling. By exploiting a defined spheroid size (d = 150 µm), the chamber is thus completely filled. Introduced spheroids subsequently merge, forming an aligned tissue, tailored by the channel geometry. (Fig. 1 B iii)

#### 3.1.3. Electrical stimulation & sensor integration

In addition to a simplified cardiac µ-tissue generation, the Spheroflow HoC directly integrates probing and analysis capabilities. It provides electrical stimulation of cultured tissues and readout of O_2_ levels via luminescent sensor spots integrated into the bottom of the tissue chamber. The integration of an O_2_ sensor spot in the knob region directly below the tissue allows for real-time measurements of O_2_ at tissue level. (Fig. 1 B iii) Both features can be combined to study the effects of pacing onto O_2_ consumption. (Fig. 1 C)

No specialized fabrication procedure is needed for adding pacing capabilities, as stimulation is carried out by harnessing fluidic media connectors as electrodes. Instead of flushing a sensor layer into the assembled chip or depositing individual spots into preassembled chip components, the tissue module is directly fabricated on a sensor substrate via micromolding of the thiolene based resin NOA 81. Patterning of NOA 81 with PDMS molds was introduced a decade ago as a simple and fast method for the generation of through-hole microfluidic stencils and its applicability in microphysiological systems has been demonstrated. [31–33] By combining a tissue layer made out of O_2_ impermeable resin with an O_2_ permeable media module made out of PDMS, we generate a precisely controlled probing environment. [34] O_2_ is introduced into the tissue chamber only via defined diffusion through the membrane, such that the sensor readout is not influenced by O_2_ diffusing from the bottom or side walls.

Developed fabrication concept involves an initial deposition of O_2_ sensor spots at defined positions via a CNC-controlled microdispenser onto adhesive treated PET-foil (). (Fig. 2 A i)

**Figure 2:**
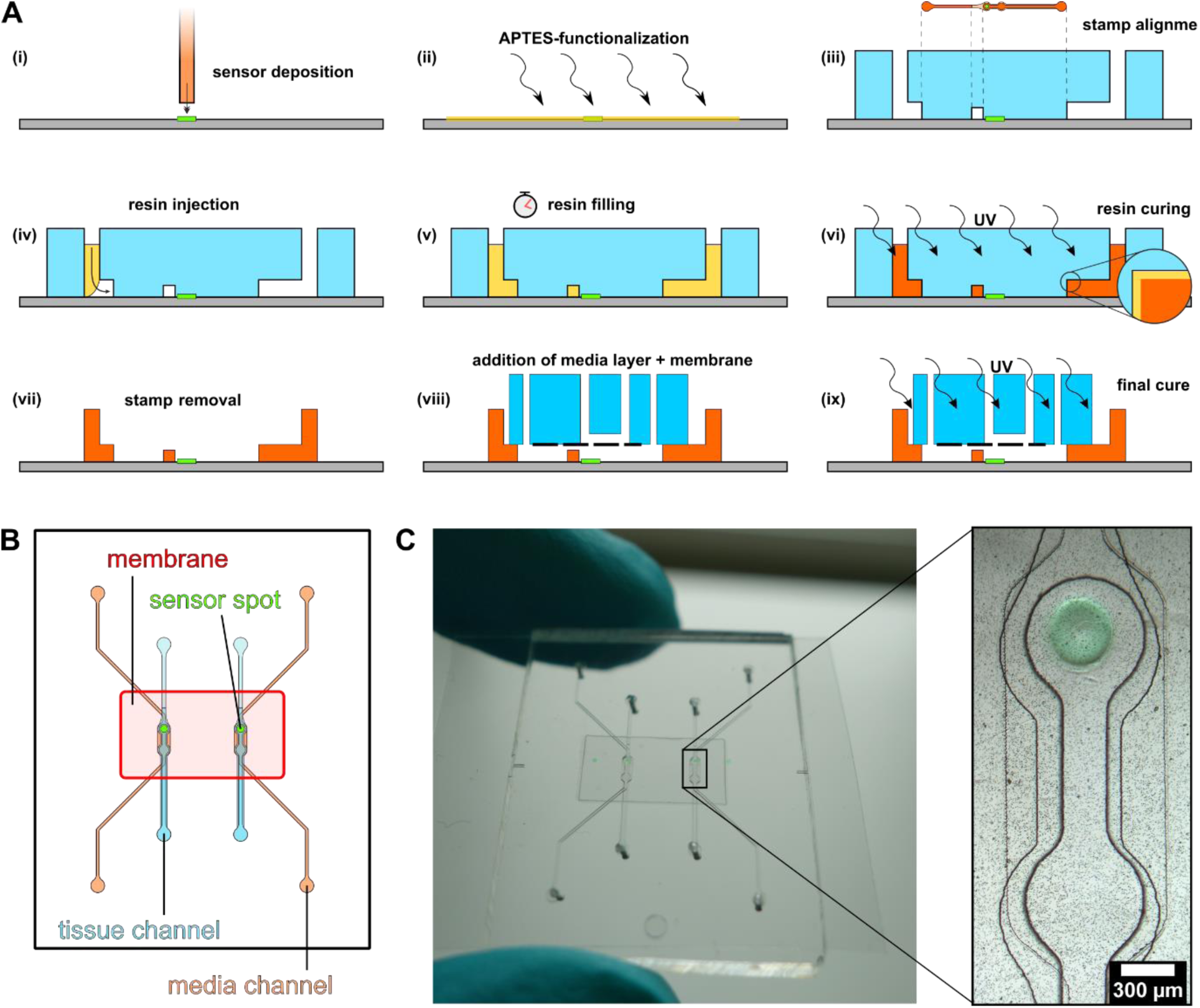
Microfluidic chip fabrication: A) Microfabrication steps (i-ix): Deposition of O_2_ sensor onto substrate (i) and further surface functionalization (ii). Addition of PDMS stamp on substrate with precise alignment of dogbone-knob onto sensor spot (iii). Injection (iv), capillary filling (v) and curing (vi) of resin. Resin in contact with PDMS does not solidify due to locally available O_2_ and remains sticky (inset). Removal of PDMS stamp exposes open tissue channels with an uncured resin layer (vii). Addition of APTES-treated media module with bonded membrane (viii) and another UV curing (ix) finalizes the chip. B) Design of Spheroflow HoC: Each chip comprises 2 individual systems consisting of a loading and media channel, separated by a porous membrane. O_2_ sensor spots are integrated into the knob region of the tissue chamber. C) Assembled chip with integrated sensors. Magnified view depicts a micrograph of a dogbone-shaped tissue chamber with integrated O_2_ sensor spot.

To ensure robust bonding between UV resin and bottom layer, the substrate is functionalized with APTES. (Fig. 2 A ii) A stamp made out of PDMS, replicating the desired microstructure of the tissue module, incorporating regions of various channel heights for channel and constriction, is placed onto the substrate. The stamp is precisely aligned, matching the second knob in flow direction of the dogbone-shaped loading chamber to the sensor spot. (Fig. 2 A iii) As the spot can be considered as flat compared to the channel structures (h < 5 µm), the elastic stamp covers the spot completely, circumventing coverage with resin in following steps. Resin is injected into the stamp injection port and completely fills the void, corresponding to prospective channel walls, within 30 min. (Fig. 2 A iv + v) The whole assembly is exposed to UV light, solidifying the bulk part of injected resin. (Fig. 2 A vi) However, resin in contact with PDMS remains uncured, as the O_2_ permeability of PDMS leads to localized availability of O_2_, preventing full curing in the contact region. After removal of the stamp, tissue channel geometries are thus replicated with a sticky surface of uncured resin, integrating the sensor spot into the channel without any coverage. (Fig. 2 A vii) The media module consisting of media channels replicated in PDMS and a joined PET membrane is treated with APTES and aligned to the tissue channel. (Fig. 2 A viii) APTES treatment enables a thorough curing without cure inhibition and thus subsequent bonding of resin to the PDMS layer. [35] Finally, the assembly is cured with UV light. One major benefit of developed fabrication procedure is the reusability of PDMS stamps (tested for > 10 molding cycles). Implemented chip design comprises 2 independent loading channels, separated by a porous membrane from individual media supplies. One O_2_ sensor spot is integrated into the rear knob of each tissue chamber. (Fig. 2 B) Finalized microphysiological system comprises a thin bottom layer, integrated sensors, and allows fluidic connections via the PDMS top layer. (Fig. 2 C)

### 3.2. Tissue characterization

Mixtures of hiPSC-derived CMs and phDFs (3:1 cell number ratio) were centrifuged into inverted microwells and reliably aggregated within 24 h to spherical spheroids of homogeneous size (d ≈ 150 µm). (Fig. 3A) For a defined aggregate composition, CM purity was assessed after each differentiation and only batches with CM purity ≥ 80% were used. Similarly, phDFs were characterized during expansion by flow cytometry using the fibroblast marker CD90. (Fig. 3B) Spheroids could be reliably introduced into the HoC, merging within 24 h and forming a compacted, aligned tissue. (Fig. 3C, SV1 + SV2) The same loading efficiency and tissue compaction was also observed in chips fabricated with a cover slip as substrate or tissue layers out of PDMS or hot embossed PET-G.

**Figure 3:**
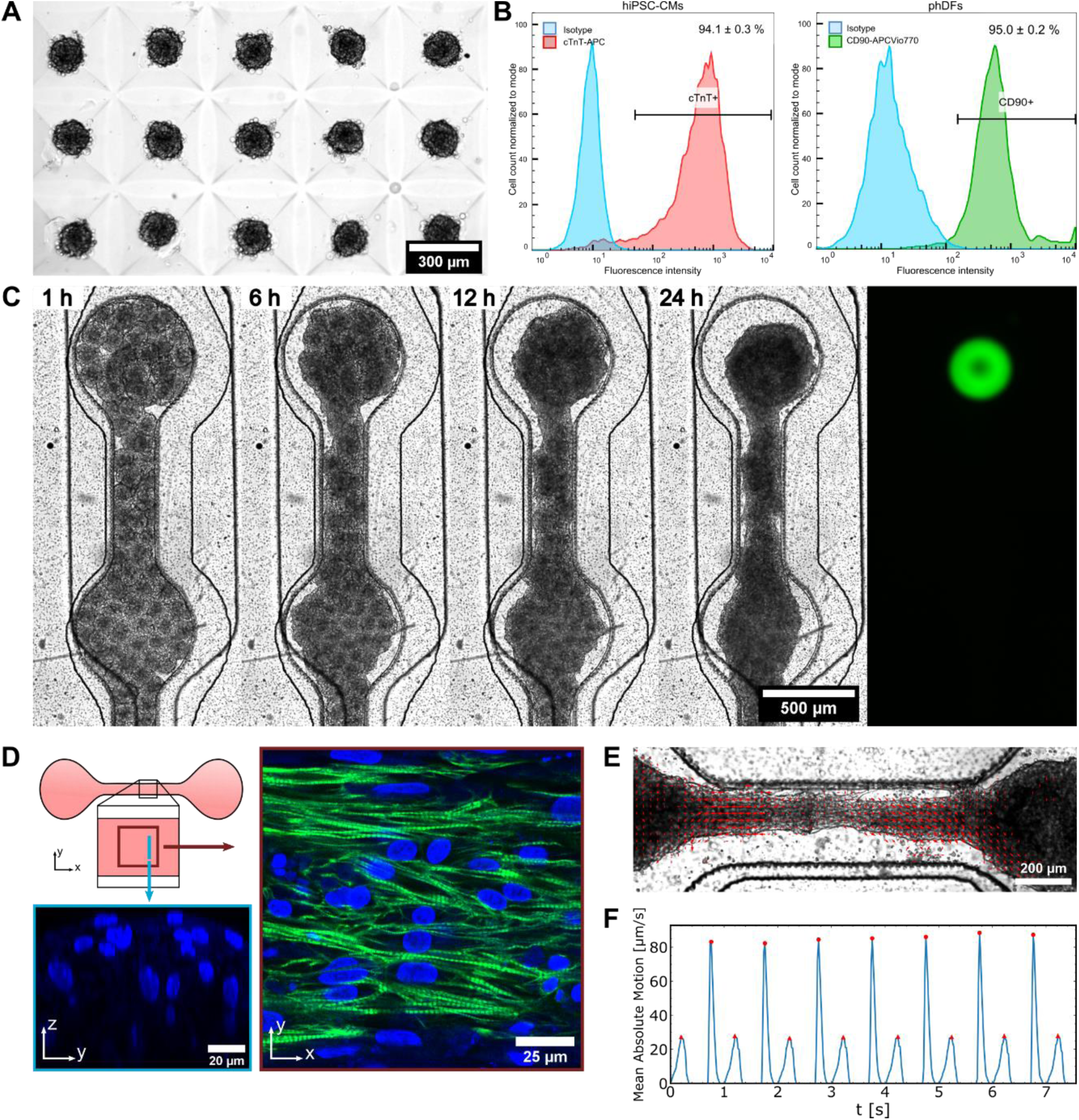
Formation of cardiac µ-tissues: A) Generation of uniformly sized spheroids in µ-Wells. B) FACS analysis of CM and FB cell populations prior spheroid formation C) Fusing of individual cardiac spheroids to an aligned tissue over the timespan of 24 h after loading. The underlying integrated O_2_ sensor spot is visualized via widefield fluorescence microscopy (green, Cy5 filterset). D) Physiological validation of generated cardiac muscle fiber in Fibronectin-coated HoC by immunostaining of cTnT (green) and DAPI (blue). The shaft region is characterized by aligned fibers (xy slice). The perpendicular z-stack view reveals stacked nuclei, indicating the generation of a multilayered 3D tissue (yz slice). E) Characterization of beating motion in cultured fiber. Motion vectors at tissue contraction reveal a collective displacement. F) Optically determined beating kinetics, amenable for beating frequency extraction.

Immunofluorescence staining revealed an alignment of cTnT fibers along the main axis of the dogbone shape. (Fig. 3D) Stacked layers of nuclei could be identified along the z-direction, verifying the generation of a 3D tissue. Hence, cells of individual spheroids reorganize, forming a single aligned cardiac fiber out of individual µ-tissues.

Usually three days after chip injection, generated tissues displayed spontaneous beating. Videos of tissue motion were recorded by video microscopy and analyzed with OpenHeartWare. The spatial distribution of tissue motion indicated a collective ordered motion, confirming the fusing of individual spheroids to a single unit. (Fig. 3E) By averaging the motion of extracted motion vector field, detailed beating kinetics could be investigated, allowing, i.a., the determination of beating frequency. (Fig. 3F)

### 3.3. Electrical stimulation of tissues

*Easypace*, an Arduino-based pulse generator, was developed to provide a cheap “DIY” platform for electrically stimulating cultured cardiac tissues. It consists of an Arduino-controlled DAC which can create two individual output waveforms, which are subsequently transformed to four independent biphasic pulses by a downstream Operational Amplifier. (Fig. 4 A) The pulse generator can be either controlled manually or remotely via Serial commands. (Fig. 4 B) Fluidic media connectors out of stainless steel (21 GA; KDS212P, Weller, USA) were harnessed as pacing electrodes and connected via alligator clips to the pulse generator. (Fig. 4 C) The theoretical electrical field distribution inside developed chip geometry was analyzed using finite elements for a voltage of U_0_ = +10 V, applied between media inlet and outlet connector. An initial study explicitly modeling individual membrane pores revealed negligible influence of introduced membrane on electrical field distribution, such that the membrane was omitted, accelerating and simplifying simulations. In the horizontal midplane, the simulation predicts a field of |E| = 0.8 V/cm that is uniform throughout the tissue chamber shaft region, which is desired for the stimulation of 3D microtissues. (Fig. 4 D)

**Figure 4:**
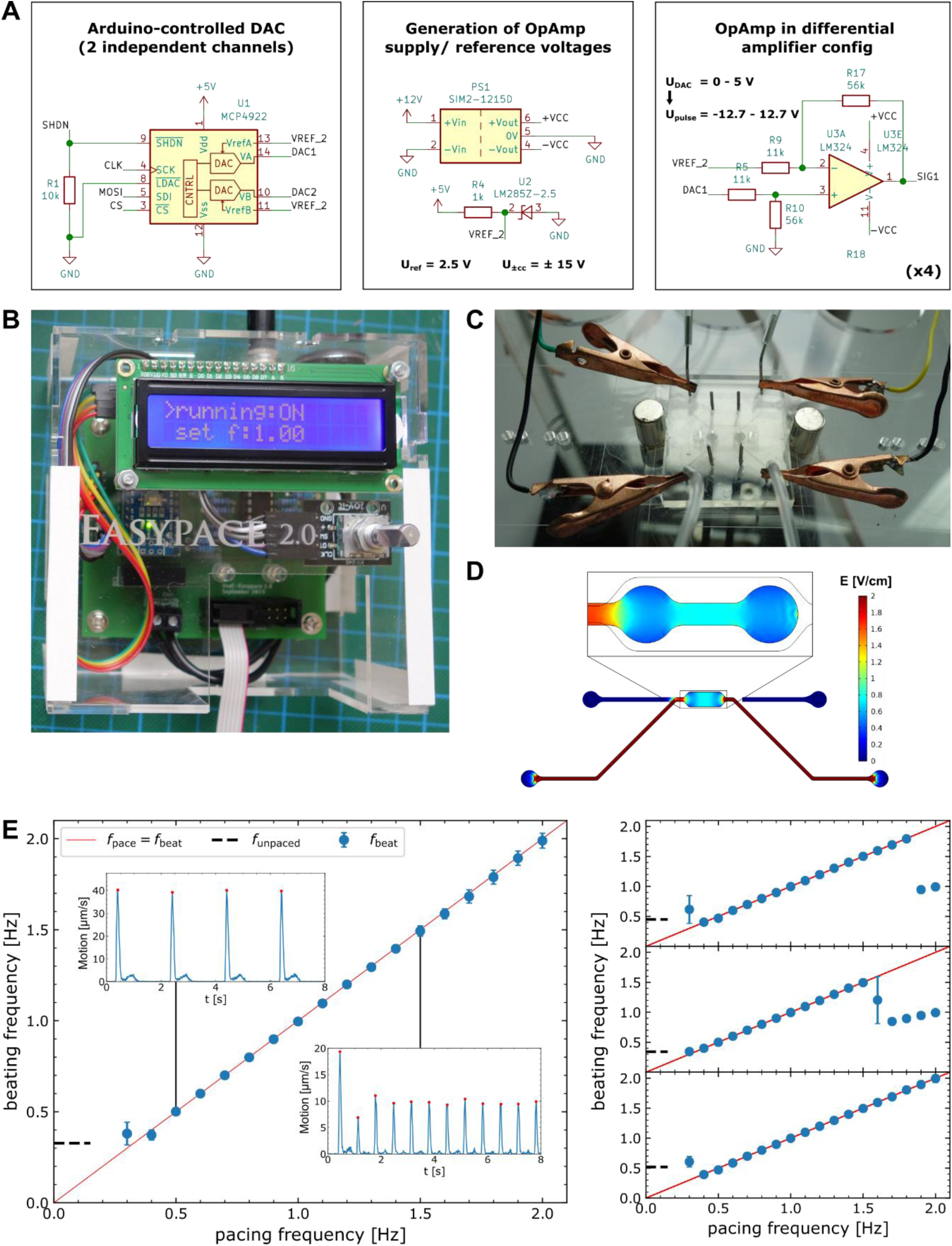
Electrical stimulation of cardiac tissues: A) Electrical schematic of key components of developed pulse generator ‘Easypace’. An Arduino-controlled DAC creates 2 independent output signals that are subsequently converted to biphasic pulses via an Operational Amplifier. B) Picture of developed pulse generator. ‘Easypace’ is controlled either manually via the rotary encoder or remotely via Serial commands and provides feedback by the integrated LCD display. C) Image of pacing setup. The pulse generator is connected via alligator clips to stainless steel fluidic connectors of the media supply, thus acting as electrodes. D) Simulated electrical field distribution inside media- & tissue-compartments upon application of 10 V between media in- & outlet. E) Beating rate analysis of tissues stimulated electrically at increasing frequencies. Beating frequency extracted from beating kinetics (insets) shown as a function of ramped up pacing frequency for four tissues. Ideal matching of beating and pacing frequency indicated by linear relation (red line).

Tissues cultured in Spheroflow chips were field stimulated on day 5 after injection. Distinct pacing frequencies ranging from 0.3 – 2.0 Hz were applied for 10 s per individual frequency and stepwise increased by 0.1 Hz. Videos of beating tissues were recorded simultaneously. By matching timestamps of each pacing frequency interval with recorded video, beating kinetics were investigated for each pacing frequency. (Fig. 4 E, inset). Comparing determined beating frequency with applied pacing frequency, a linear relation up to a maximum pacing frequency is observed. (Fig 4 E) Exceeding this frequency, tissues could not follow the external pacing rate anymore. They skipped every second pulse, yielding a beating frequency half of applied frequency. All investigated tissues could be paced up to 1.5 Hz, individual tissues up to the applied maximum pacing rate of 2.0 Hz. Our findings confirm the applicability of employed facile pacing approach via fluidic media connectors. Without the integration of additional electrodes, tissues can be paced and the beating rate precisely controlled.

### 3.4. Investigation of O_2_ levels under varying metabolic activity

To assess the functionality of integrated O_2_ sensing spots, sensor spots in chips with cultured beating cardiac tissues were monitored overnight. Cultured tissues were electrically stimulated by applying a cyclical variation of consecutive biphasic pulses of frequencies 0.7, 1.0, 1.2, and 1.5 Hz. Stimulation was carried out for each frequency for 10 Minutes with a resting phase of 30 Minutes between the application of successive frequency. (Fig. 5 A) Each frequency cycle was repeated for three times. In order to exclude any effects of external influences on O_2_ levels, pacing cycles were temporally shifted for both paced systems by 20 Min with respect to each other. Upon initiation of pacing, a drop in measured O_2_ levels can be distinguished, with drop onset clearly coinciding with pacing start. (cf. reference line) As in the unpaced reference tissue constant O_2_ levels are detected, this drop can be attributed to a higher tissue O_2_ consumption that might be caused by increased metabolic activity, triggered by electrical stimulation. Upon termination of pacing, an increase in O_2_ levels towards initial partial pressures, monitored in the unpaced state, is observed. Stimulating the other tissue, the identical behavior is detected. Increasing pacing frequencies display increased jumps in O_2_ levels. Jump height is determined by the difference between O_2_ partial pressure at pacing onset and pacing termination, and evaluated for both tissues as a function of applied pacing frequency. Jump heights are furthermore normalized to the difference of O_2_ partial pressure between the unpaced state and the fully oxygenated state, yielding relative jumps in O_2_ concentration. (Fig. 5 B) Relative jumps expose for both investigated systems an increase with applied pacing frequency, observable in each pacing cycle.

**Figure 5:**
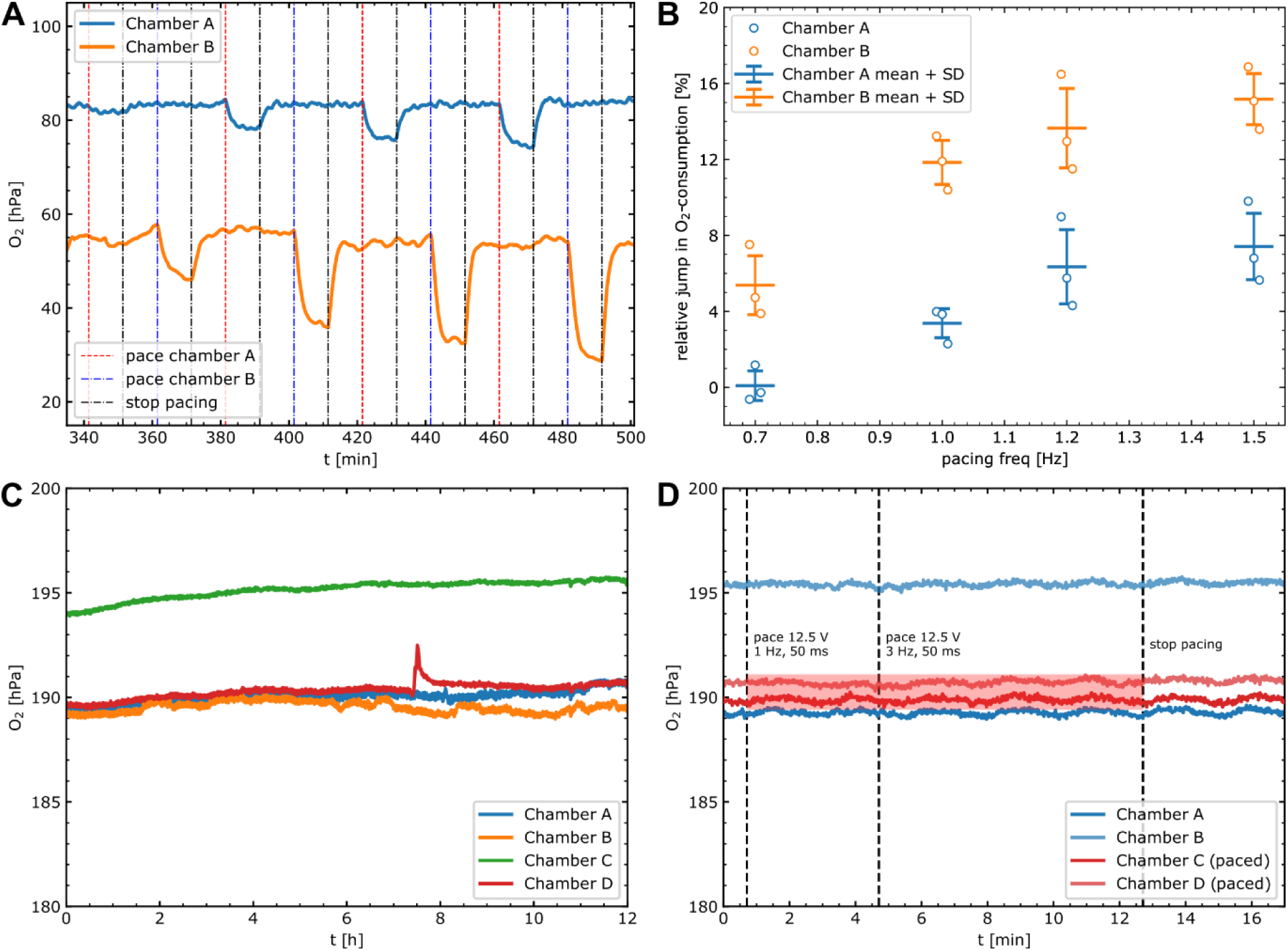
Integrated measurements of O_2_ levels: A) Recording of O_2_ partial pressures of 2 tissues alternatingly paced (dashed lines) at increasing frequencies of 0.7, 1.0, 1.2, 1.5 Hz. B) Evaluation of frequency dependence of relative jump heights in O_2_ consumption for tissues repeatedly probed following timings in (A). C) Stability analysis of O_2_ levels in empty systems perfused with media over 12 h. D) Comparison of O_2_ partial pressures between paced and unpaced chambers, measured in empty, media-perfused systems. No difference between both conditions is observed, excluding O_2_ changes induced by electrolysis.

Signal stability of developed sensor chip is investigated by perfusing four empty systems with media in culture conditions (37 °C, 5 % CO_2_, 50 µl/h). Over the timespan of 12 h, all measured O_2_ partial pressures display minimal (< 1 hPa/h) shifts, precluding O_2_ scavenging of employed resin NOA 81. (Fig. 5 C) Chamber 4 exhibits a spike in O_2_ partial pressure, which we attribute to a passing air bubble.

In order to attribute the observed variation in O_2_ levels to an increase in tissue O_2_ consumption, excluding potential changes in O_2_ concentration triggered by electrolysis, O_2_ levels were recorded in an empty chip perfused with media, undergoing electrical stimulation. (Fig. 5 D) Two out of four investigated systems were stimulated at the maximum voltage of 12.5 V with initial frequency of 1 Hz, further ramped up to 3 Hz for up to 8 min. No drop in O_2_ concentration is detected. Comparing the O_2_ partial pressure trend with unpaced chambers, all exhibit a similarly constant trend.

## 4. Discussion

Many concepts of HoCs have been proposed in recent years, in this work we have presented a novel microphysiological system targeting controlled tissue generation, furthermore, directly integrating O_2_ sensing and electrical pacing capabilities. Demonstrated tissue generation approach combined the well-established technique of cardiac spheroid formation with OOC technology. Thus, aligned cardiac fibers could be generated out of radially symmetric multicellular cardiac spheroids of defined composition as starting point. Developed bottom-up approach of preforming spheroids with subsequent merging provided robust loading and standardized µ-tissue generation. Due to the simple loading mechanism, driven by hydrostatic pressure, presented system offers opportunities for an automatized tissue generation, feasible even at large-scale, as only simple pipetting steps are necessary which could be carried out by liquid handling robots. Currently, each injection port is connected to only one chamber, a conceivable multiple branching of the filling channel however offers potential for parallelization.

Compared to single cells, which can hardly be compacted in a tissue chamber without applying uncontrolled pressures, the utilization of spheroids as building blocks facilitates injection as they can easily be guided by a clogging-free side lying constricted channel to desired positions. We presented a dog bone shaped tissue chamber, however the underlying concept is universal, allowing µ-tissue accumulation in arbitrary formed tissue chambers or even the introduction of other cell types. As spheroids fill the tissue chamber subsequently, it is also conceivable to consecutively inject spheroids of varying composition, thus creating, e.g., a cardiac fiber which couples fibrotic tissue on one end with healthy tissue on the other end. Furthermore, proposed concept is not fixed on the utilization of a sensor substrate, we assembled chips of the same design on coverslips for high-resolution imaging or fabricated a pure PDMS based system integrating two micropillars inside the dog bone knobs enabling readouts of contractile force in further studies. Utilized PET substrate was chosen to provide optimal sensor spot adhesion, however we observed corrupted imaging qualities using epi-illumination microscopy. Thus, the media layer had to be removed for immunofluorescence imaging. We plan to exchange utilized substrate in further chip iterations, enabling imaging of the chip in the unperturbed state while maintaining sufficient sensor spot adhesion.

We demonstrated a noninvasive integration strategy of electrical pacing capabilities, superseding complex integration of electrodes into the chip for simple pacing purposes, and could verify precise control of beating rate with chosen approach. Pacing was only investigated for short time periods (up to 10 Min); long-term pacing with applied pacing voltage might lead to the creation of toxic byproducts. However, compared to pacing in a static system, we are confident that existing media flow removes toxic byproducts, such that pacing can be also carried out for longer time periods, e.g. for inducing maturation.

We are aware that predicted field strengths of 0.8 V/cm lie below field strengths often used in literature of 5 V/cm. [36] Utilized parameters however still induced robust pacing which is most probably balanced by an increased pulse width of 50 ms compared to commonly used 5 ms. If higher fields are desired, utilizing tissue channel sealing plugs as electrodes is also feasible, allowing in addition a flexible choice of electrode material.

Within our studies we developed the low-cost pulse generator *Easypace*, which we think is a valuable Open-Source Hardware tool for the scientific community with its easy fabrication and scriptable pacing capabilities. Provided automation capabilities are of special interest for running overnight experiments involving various pacing parameters or frequency sweeps.

Standardization and upscaling of existing OOC platforms is essential for any application in an industrial setting. Developed novel sensor integration concept is distinguished by its approach of directly fabricating the chip on top of the sensor substrate. Hence, it is not necessary to ship pre-assembled chip components to external facilities for sensor deposition. Standardized sensor substrates, offering a geometrical arrangement of sensor spots on a predefined regular grid, which are not tailored to any specific chip design, could be produced in large quantities in the future and lead to a broader adaption of the integration of sensors into OOC platforms due to wide availability, furthermore guiding chip design.

By integrating the O_2_ sensor spot into the bottom of the tissue chamber, the sensor is in direct contact with cultured tissue, granting *in situ* readouts and minimizing aberrations from spatial oxygen gradients. As employed UV resin is gas-impermeable, a defined system with O_2_ influx only via the membrane is generated.

We verified stable O_2_ measurements without O_2_ scavenging of employed UV resin. Comparing signals of various chambers perfused under identical conditions, we noticed slight differences in measured O_2_ partial pressures (up to 5 hPa). We attribute these deviations to variations in the manual fabrication process. Slight variations in stamp alignment can lead to partial coverage of the sensor layer with resin. As the covered part is not aerated, this might lead to a biased signal. The same calibration was used for all sensor spots, not accounting for individual variations. We suggest an individual calibration of each sensor in further studies quantifying O_2_ consumption rates.

Ultimately, we provided a biological proof-of-concept of combined stimulation and probing inside a HoC, assessing metabolic tissue activity under the influence of external pacing. Integrated O_2_ sensors granting fast *in situ* read outs offer huge potential to assess changes in metabolism triggered by other biochemical or mechanical stimuli. Particularly, we see huge potential for an integrated analysis of oxygen consumption rates in cultured µ-tissues, similarly to the Seahorse assay, the gold standard for metabolic measurements. In addition to O_2_ sensing, developed platform can be used to study Ca propagation kinetics along the uniaxial cardiac fiber. (cf. supplementary) We determined mean Ca propagation velocities within a fiber cultured in the chip to v_sig_ = 55 mm/s, coinciding in magnitude with velocities measured in similar systems of 46 mm/s and 95 mm/s. [37,38]

All in all, developed system can be considered a starting point opening the door for addressing key biological questions, e.g., investigating maturation via pacing or the effect of administered drugs on O_2_ consumption of physiological µ-tissues.

## Data availability

All experimental data within the article are available from the corresponding author upon reasonable request.

## Credit author statement

Oliver Schneider: Conceptualization, Methodology, Investigation, Software, Data curation, Visualization, Writing-Original draft preparation; Alessia Moruzzi: Methodology, Investigation, Writing-Original draft preparation, Writing-Reviewing and Editing. Stefanie Fuchs: Conceptualization, Methodology; Alina Grobel and Henrike S. Schulze: Investigation; Torsten Mayr: Funding acquisition, Supervision; Peter Loskill: Writing - Review & Editing, Funding acquisition, Supervision.

## Declaration of competing interest

The authors declare the following competing interest(s): T.M. is a founder, holds equity in PyroScience GmbH in Germany, and is the CEO of the Austrian branch, PyroScience AT GmbH. PyroScience is a developer, producer, and vendor of sensor technology.

## Acknowledgments

The authors would like to thank Sarah J. Rockwood, Ana C. Silva and Todd C. McDevitt at Gladstone Institutes for an introduction and training on spheroid formation and culture.

The research was supported in part by the DAAD funded by the Bundesministeriums für Bildung und Forschung (BMBF) (PPP USA 2018, 57387214) as well as the European Union’s Horizon 2020 research and innovation program under the Marie Skłodowska-Curie grant agreement no. 812954.

## Supplementary information

**Figure S1:**
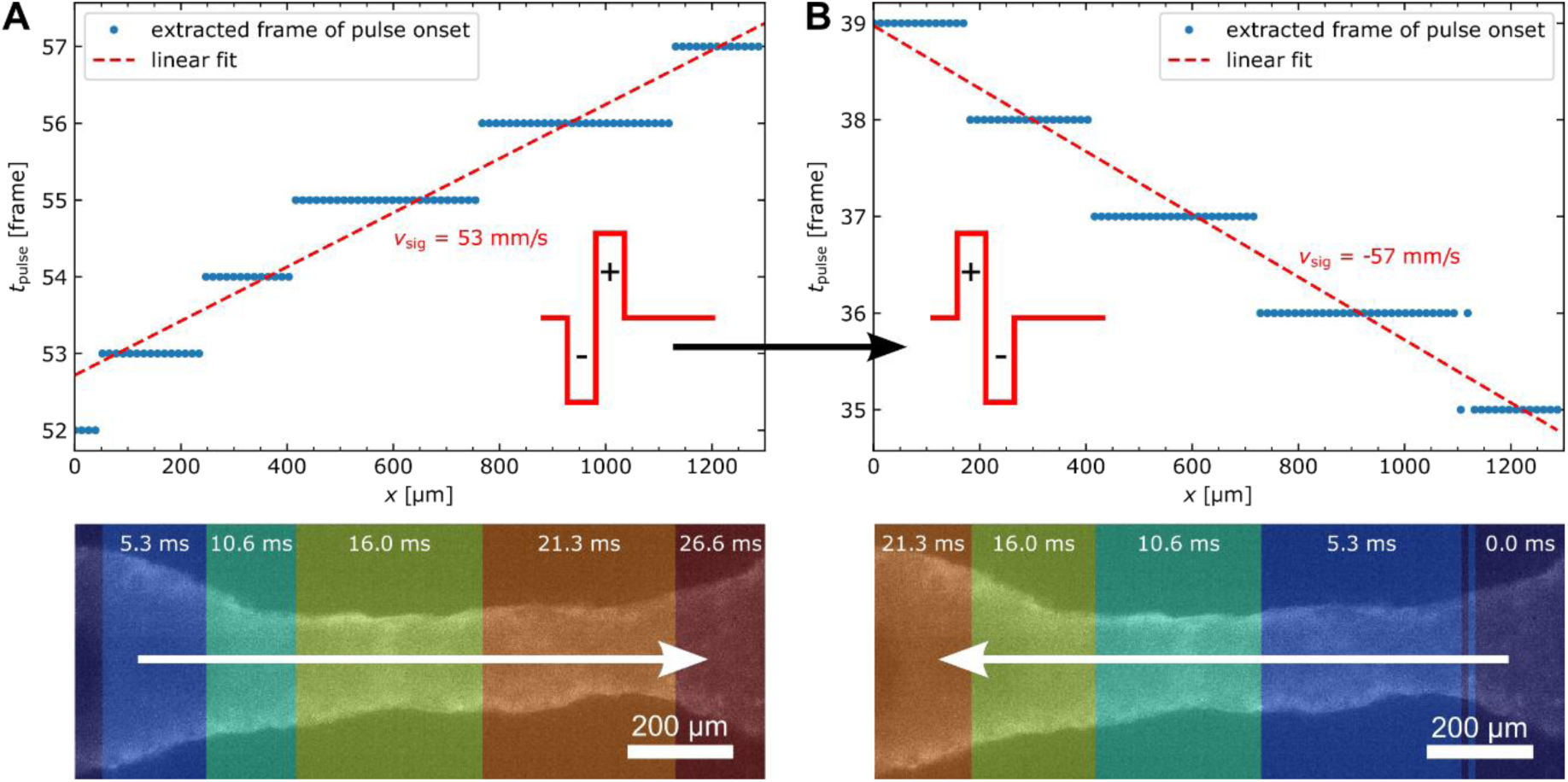
Analysis of Ca propagation velocity: A) Evaluation of pulse onset of paced tissue as a function of fiber position, extracted from high-speed video microscopy recording of Fluo-4 AM Ca-dye. Ca propagation velocity v_sig_ is determined from linear fit. Color-coded Ca activation map is shown as overlay to the investigated tissue. B) Analysis of pulse onset for inverted pacing polarity. Propagation velocity is determined and propagation shown as overlay similar to (A).

### Ca analysis

Propagation of Ca signaling within cultured tissues was analysed by fluorescence video microscopy. Media supply tubing was disconnected and the media channel flushed with dye solution (10 µM Fluo-4 AM [F14201, Invitrogen, USA] in Tyrode’s solution [T2397, Sigma-Aldrich, USA]) via pipet tips by inserting an empty tip into the outlet, adding a tip filled with 200 µl dye solution into the inlet, and starting the flow by manually applying slight pressure. The chip was subsequently incubated in the dark for 15 min and transferred to the microscope with integrated incubator. Already inserted stainless steel plugs sealing the tissue compartment were utilized as pacing electrodes and connected with alligator clips to the pulse generator. Tissues were paced at 1 Hz with positive as well as negative polarity of the first pulse phase and the fluorescence signal of the Ca dye recorded at 188 fps. Cultured cardiac fiber was aligned parallel to the image x-axis during recordings. Recorded signal was averaged for each frame along the y-axis and binned in 10 pixel bins along the x-axis yielding a 1D intensity distribution for each timepoint. For each position on the x-axis, the frame of signal onset was determined by the videoframe in which the maximum of signal derivative occurred. From a linear fit of activation time vs. activation position, the signal propagation velocity can be determined to v_sig_ = 53 mm/s and v_sig_ = -57 mm/s for positive and negative pulse polarities. An inversion of signal propagation direction is monitored upon pacing polarity inversion.

### Time-lapse of tissue formation

Following spheroids injection, pipet tips in tissue inlet & outlet were exchanged to filter tips containing fresh prewarmed media (100 µl per tip) and chips fixated on the motorized stage of an optical microscope with integrated incubator (37 °C; DMi8,Leica, Germany). Tissue formation in microphysiological systems loaded with CMs (SV1) as well as FBs (SV2) was monitored by acquiring images in 30 min intervals. For both tissue types merging of individual spheroids as well as tissue compaction is observed.

### Pacer assembly

*Easypace* can be beginner-friendly assembled from broadly available, hand-solderable parts. The core component is a custom pcb, which can be ordered from a variety of suppliers. It is equipped with an Arduino Nano for coordinating user input and pacing cycles. Detailed build instructions, a bill of materials as well as example pacing scripts for controlling *Easypace* remotely can be found in the Github repository. (https://github.com/loslab/easypace/)

### Dye dispensing parameter

**Table T1:**
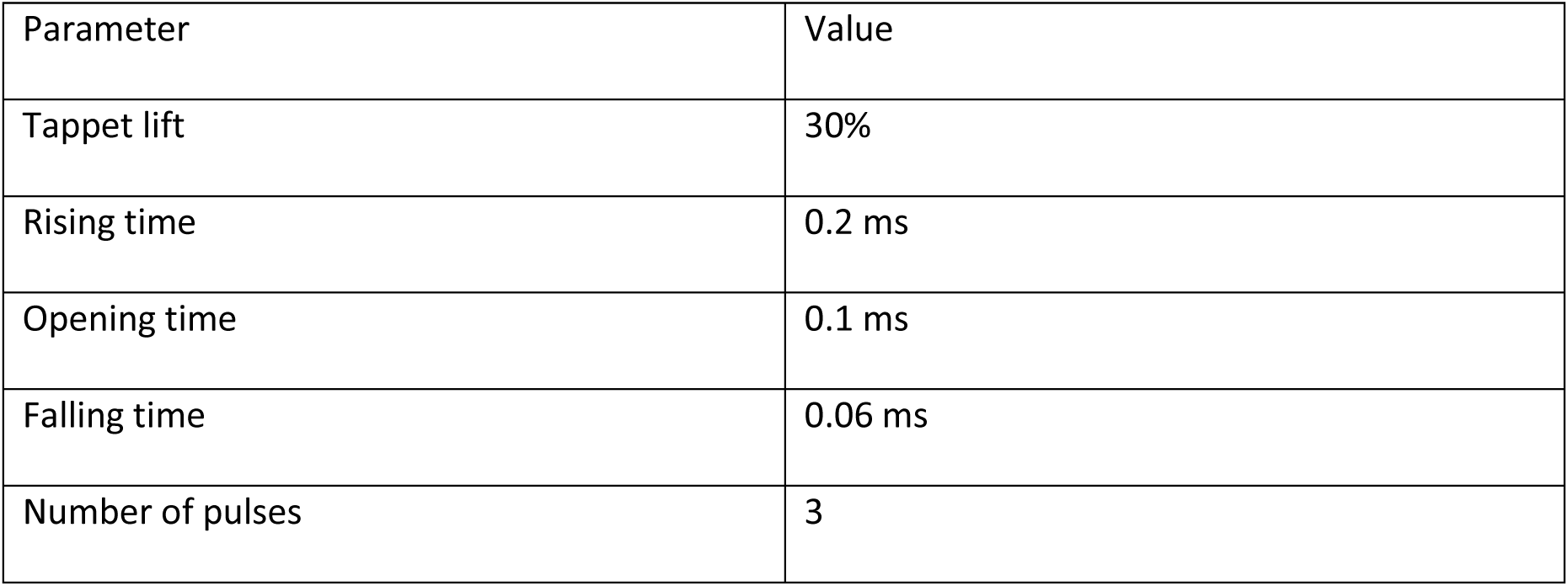
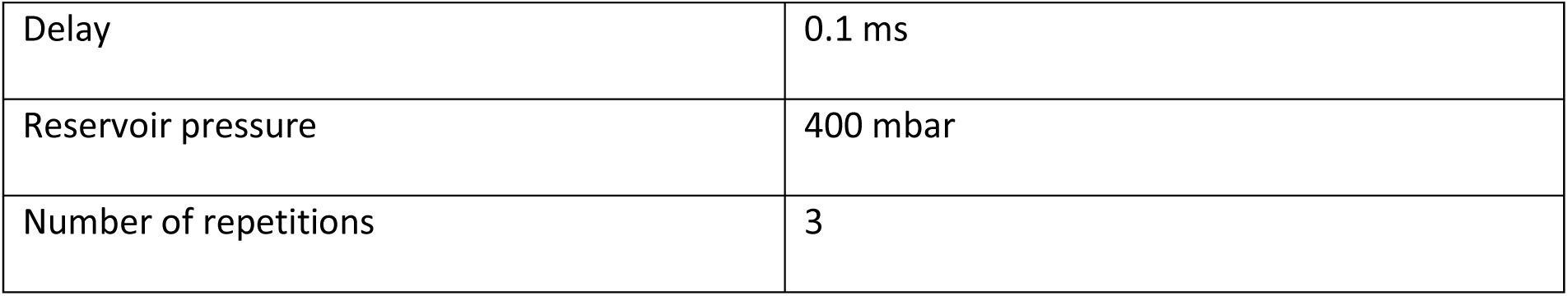
Microdispensing parameter for deposition of luminescent O_2_ dyes

## Notes

### Competing Interest Statement

The authors declare the following financial interests/personal relationships which may be considered as potential competing interests:
Torsten Mayr reports a relationship with PyroScience GmbH that includes: equity or stocks. The authors declare the following competing interest(s): T.M. is a founder, holds equity in PyroScience GmbH in Germany, and is the CEO of the Austrian branch, PyroScience AT GmbH. PyroScience is a developer, producer, and vendor of sensor technology.

